# Carbonized rubber electrodes can cause a DC-offset in transcranial alternating current stimulation

**DOI:** 10.1101/2025.09.11.675526

**Authors:** Heiko I. Stecher, Peppi Schulz, Arnd Meiser, Alexander Hunold, Klaus Schellhorn, Christoph S. Herrmann

**Affiliations:** Experimental Psychology Lab, Department of Psychology, Carl-von-Ossietzky Universität Oldenburg, Ammerlander Heerstr. 114-118, 26111, Oldenburg, Germany; European Medical School, Cluster for Excellence “Hearing for All”, Research Center Neurosensory Science, Carl-von-Ossietzky Universität Oldenburg, Carl-von-Ossietzky-Straße 9-11, 26129, Oldenburg, Germany; Research Center Neurosensory Science, Carl von Ossietzky Universität, 26129, Oldenburg, Germany; Institute of Biomedical Engineering and Informatics, Technische Universität Illmenau, Gustav-Kirchhoff Str. 2, 98693 Ilmenau, Germany; neuroConn GmbH, Ilmenau, Germany

**Author notes:** Corresponding author: Christoph S. Herrmann.

## Abstract

**Introduction:** Carbonized rubber electrodes are widely used in non-invasive brain stimulation studies. Due to their polarizable nature, however, they can cause a voltage offset, which might be problematic for concurrent EEG studies.

**Objective:** In this study, we aim to describe the voltage offset and ensure that the offset does not alter the intended waveform of applied stimulation.

**Methods:** Using data from 2 human studies and phantom measurements, which employed carbonized rubber electrodes, we quantify the magnitude and frequency of DC-offsets and contrast this against pilot-measurements using Ag/AGCl-electrodes. In a further phantom study, we record the offset-voltage that arises from the electrode/electrolyte interface and compare this to the voltage put out by the stimulation device.

**Results:** A non-zero voltage offset is present in all human and phantom studies employing carbonized rubber electrodes, while the offset using Ag/AgCl electrodes is close to zero. Direct measurements of the stimulator output in the presence of a measurable voltage offset at the stimulation electrodes shows that the offset originates from the electrodes and not from the current provided by the stimulation device.

**Conclusion:** Using carbonized rubber-electrodes for stimulation can result in the emergence of a measurable voltage offset, due to their polarizable nature. We argue that this offset can be problematic in concurrent EEG recordings, as they pose the risk of amplifier saturation and distortions of the recorded stimulation waveform.

---

Non-invasive brain stimulation methods such as transcranial electrical stimulation (TES) are highly valuable tools for investigating the causal neurophysiological underpinnings of human behavior (Herrmann et al., 2013) and hold promise for therapeutic applications (Elyamany et al., 2021). In transcranial alternating current stimulation (tACS), a sinusoidal current oscillating around zero without a DC-component (offset) is applied. In our lab, this is achieved by generating a digital sine-wave, which is delivered via a digital to analog converter to the stimulation device (DC-STIMULATOR PLUS, neuroConn, Ilmenau, Germany) operating in remote mode. The stimulator converts this input voltage to a constant current. During no-stimulation blocks, a 0 V input is applied, resulting in a 0 mA output current.

However, when employing carbonized rubber electrodes with conductive paste (ten20, D.O. Weaver, Aurora, CO, United States), as is common practice in TES studies (Woods et al., 2016), we observed that our stimulation devices occasionally report a gradually increasing, non-static offset-voltage even when the input signal remains at 0 V.

In two pre-registered concurrent EEG/alpha-tACS experiments from our lab (unpublished data), we manually recorded the electrode voltage as given by the device in remote mode during no-stimulation blocks. The pre-registrations for Experiment 1 (N = 32) and Experiment 2 (N = 20) can be found in https://osf.io/qw286 and https://osf.io/85kb7. A brief description of both experiments is provided in the supplementary material.

In the verum conditions, we observed an increase in the average absolute offset from beginning to the end of the experiments. Comparable offsets were also recorded during sham conditions, suggesting independence from stimulation type. An overview of offset voltages and impedance values is given in Figure 1 A (left) and in Supplementary Tables 1 and 2.

**Figure 1:**
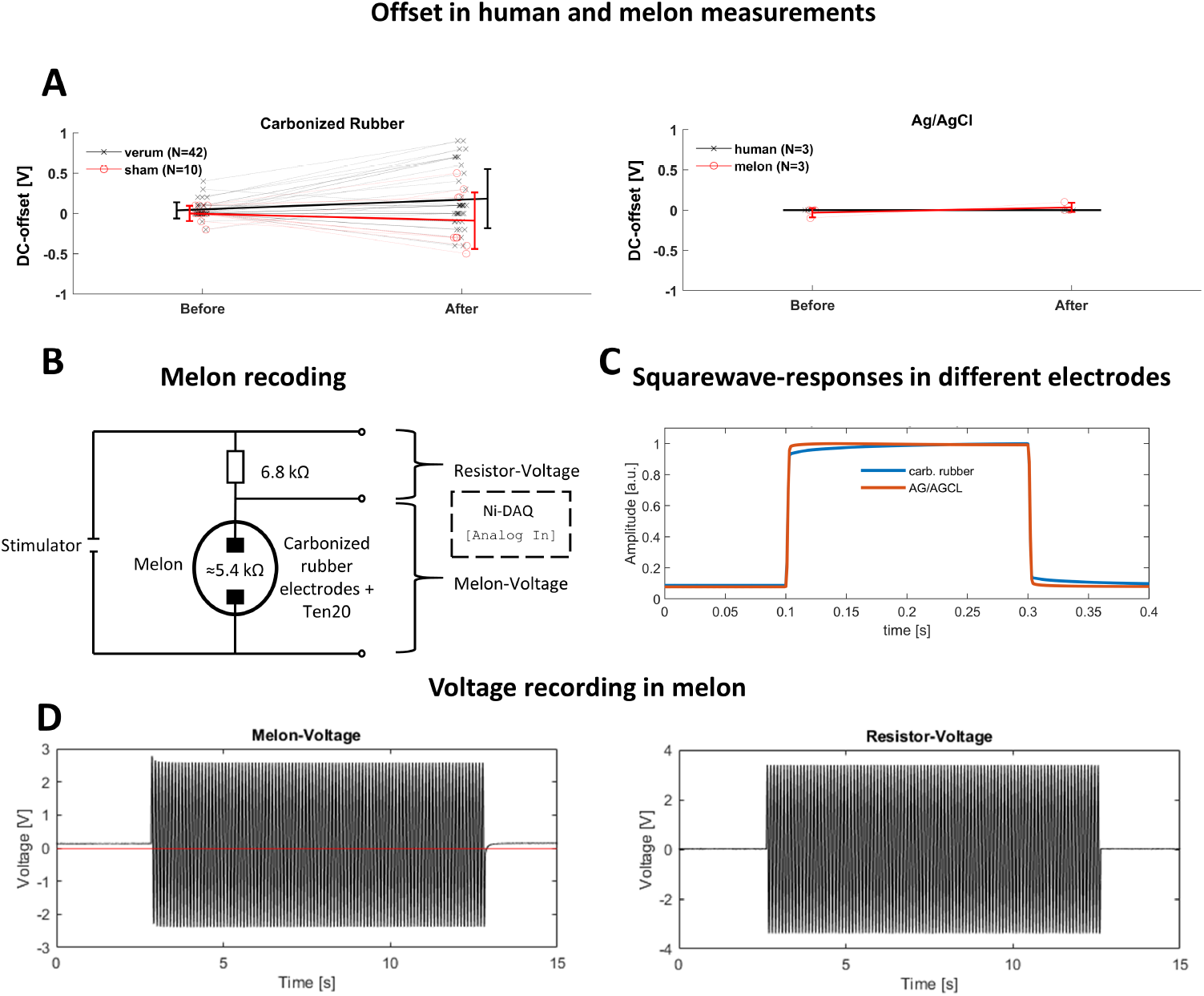
**A:** *Recorded offset before and after 21 minute tACS in humans and melons:* Left using carbonized rubber, right using Ag/AgCl-electrodes. Solid lines depict mean and standard deviation. **B:** *Setup to record DC-offsets in a melon:* Carbonized rubber stimulation electrodes were affixed on a watermelon roughly 10 cm apart with Ten20 conductive paste. The impedance was brought to ≈ 5.5 kΩ and the electrodes connected to a Stimulator DC Plus. For the cathode, a 6.8 kΩ resistor was placed between stimulator and electrode. Voltage at the resistor and between the electrodes were recorded by using a NI USB 6251 (National Instruments, USA) A/D converter. **C:** *EEG-recordings of square-wave stimulation pulses:* Each recording shows mean of 60 trials, using either rubber electrodes (blue), or Ag/AgCl-electrodes (red). **D:** *A/D-converter recording of stimulation signal:* 10 s, 10 Hz, 0.5 mA (peak to peak)-sine-wave at phantom melon. Left: recorded from electrodes placed on the melon.

**Figure 2:**
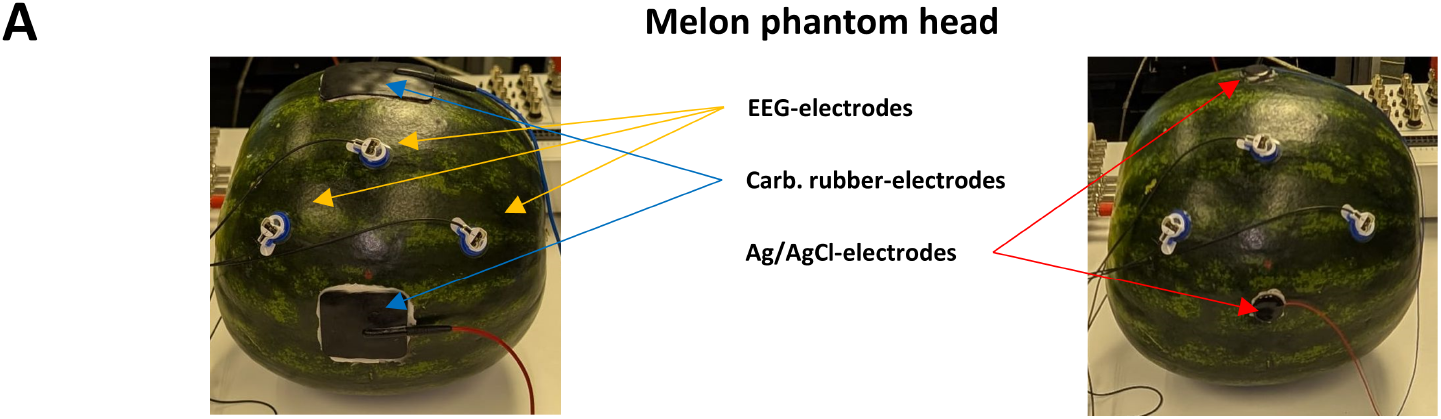
**A:** Exemplary setup of a melon phantom head to record offsets. Three active EEG electrodes were placed at positions roughly corresponding to P3, Pz and P4 (REF on nose, GND on FPz). Left: Carbonized rubber stimulation electrodes were placed at Cz and Oz. Right: At the same position Ag/AgCl-electrodes for stimulation were placed. Right: recorded from resistor placed between stimulator and electrode. Sold red line in left depicts 0 V.

Phantom measurements with a watermelon (*Citrullus lanatus*) yielded similar offset values (e.g. 0.9 V offset after 5 minutes of tACS at 5 kΩ impedance; see Supplementary Table 1).

These DC-offsets are difficult to detect in concurrent EEG recordings as they cannot be distinguished from naturally occurring drifts. However, they can be measured directly using multimeters, oscilloscopes, or A/D-converters as a voltage between the electrodes (see Fig. 1, B and D, left. Notably, the voltage before and after stimulation deviates from zero (red line) at ≈ 0.2 V, consistent with the voltage as reported by the stimulator. Wafeform distortions were also observed at the onset and offset of stimulation. Importantly, these voltages could only be recorded from the melon and not from a resistor placed in sequence with the melon inside the stimulation circuit (see Fig. 1B and 1D, right), indicating that the potential does not stem from a true DC current generated by the stimulator.

We attribute these DC-offsets to the polarizable nature of carbonized rubber electrodes (Agrebi et al., 2017; Zhu & Deegan, 2024). Supporting this interpretation, Fig. 1 C displays distortions of perfect square waves by the rubber-electrodes. We propose that the electrode-electrolyte-interface becomes charged for instance through leakage currents and/or asymmetrical currents during impedance checks.

Many researchers record EEG during transcranial brain stimulation, either to capture the stimulation signal for subsequent artifact correction in tACS, or to obtain artifact-free recordings by the use of transcranial temporal interference stimulation (von Conta et al., 2022) and transcranial amplitude modulated alternating current stimulation (Witkowski et al., 2016). In this context, we argue that polarizable electrodes pose two problems: 1) DC-offset may be added to the cortical EEG-signals, risking driving the EEG-amplifier saturation and 2) non-linear transfer-functions may distort the recorded waveform, producing artifacts at envelope frequencies.

Ag/AgCl-electrodes, considered non-polarizable, may provide a viable alternative. Preliminary experiments from our lab in both humans (n=3) and melons (n=3), using Ag/AgCl-electrodes attached with the same conductive paste and stimulation devices, showed a reduction of the DC-offset to *<*0.1 V after 21 minutes of alternating current stimulation (see Fig. 1A, right).

Nevertheless, potential drawbacks of Ag/AgCl electrodes, include higher costs, and limited lifespan. Ag/AgCl electrode degradation – probably due to electrolysis – has been reported after 8 sessions of tDCS, leading to increased impedances and unpredictable changes in the applied electric field (Langenbach et al., 2020). Hampstead et al. (2020) suggest that proper rotation of electrodes used as anodes or cathodes could slightly increase their lifetime to up to ~ 10 sessions. To date, no clear recommendations exist for tACS. It is conceivable, however, that electrode degradation electrodes is less pronounced in tACS, since electrolysis requires direct currents. Until further guidelines are available, we recommend careful inspection of Ag/AgCl for degradation before each session.

In summary, rubber-electrodes can become polarized, during concurrent EEG recordings the resulting DC offset of up to ?*sim*1*V* may saturate the amplifier. To address this, researchers may opt for non-polarizable stimulation electrodes or apply hihgh-pass filters to remove DC components from EEG recordings.

## CRediT authorship contribution statement

Heiko I. Stecher: Writing – original draft, Visualization, Software, Methodology, Investigation, Formal analysis, Data curation, Conceptualization. Peppi Schulz: Writing – original draft, Software, Methodology, Formal analysis, Data curation, Conceptualization. Arnd Meiser: Writing – review & editing, Methodology, Investigation, Conceptualization. Alexander Hunold: Writing – review & editing, Methodology, Conceptualization. Klaus Schellhorn: Writing – review & editing, Methodology, Conceptualization, Supervision. Christoph S. Herrmann: Writing – review & editing, Writing – original draft, Supervision, Methodology, Investigation, Funding acquisition, Conceptualization.

## Supplementary Material

**Table 1:**
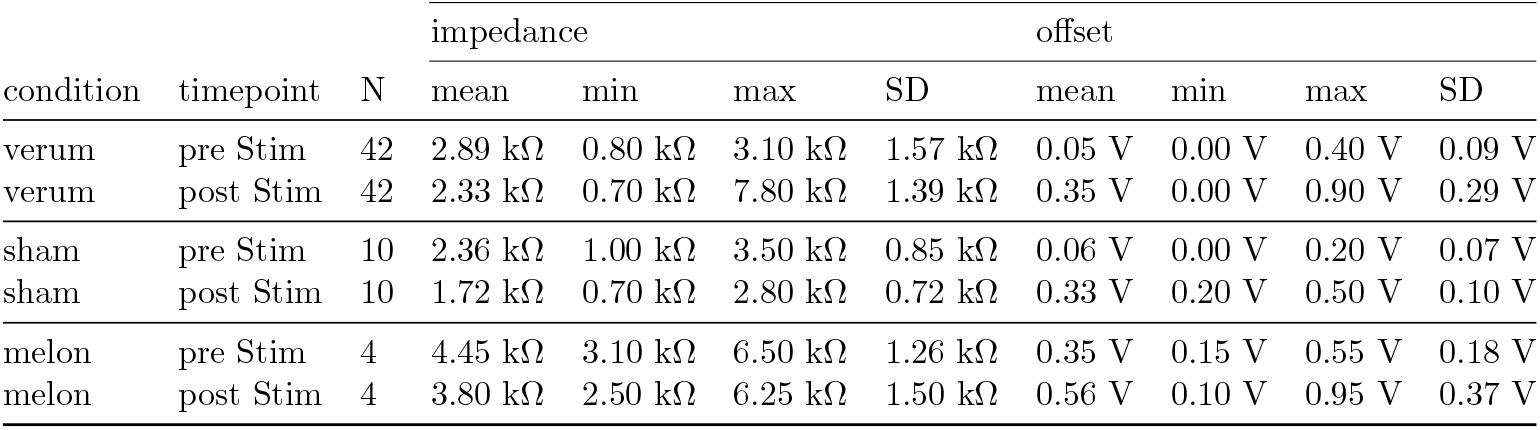
impedances and absolute offset voltages recorded in our tACS experiments using rubber electrodes.

| condition | timepoint | N | impedance |  |  |  | offset |  |  |  |
| --- | --- | --- | --- | --- | --- | --- | --- | --- | --- | --- |
|  |  |  | mean | min | max | SD | mean | min | max | SD |
| verum | pre Stim | 42 | 2.89 kΩ | 0.80 kΩ | 3.10 kΩ | 1.57 kΩ | 0.05 V | 0.00 V | 0.40 V | 0.09 V |
| verum | post Stim | 42 | 2.33 kΩ | 0.70 kΩ | 7.80 kΩ | 1.39 kΩ | 0.35 V | 0.00 V | 0.90 V | 0.29 V |
| sham | pre Stim | 10 | 2.36 kΩ | 1.00 kΩ | 3.50 kΩ | 0.85 kΩ | 0.06 V | 0.00 V | 0.20 V | 0.07 V |
| sham | post Stim | 10 | 1.72 kΩ | 0.70 kΩ | 2.80 kΩ | 0.72 kΩ | 0.33 V | 0.20 V | 0.50 V | 0.10 V |
| melon | pre Stim | 4 | 4.45 kΩ | 3.10 kΩ | 6.50 kΩ | 1.26 kΩ | 0.35 V | 0.15 V | 0.55 V | 0.18 V |
| melon | post Stim | 4 | 3.80 kΩ | 2.50 kΩ | 6.25 kΩ | 1.50 kΩ | 0.56 V | 0.10 V | 0.95 V | 0.37 V |

**Table 2:** impedances and absolute offset voltages recorded in our tACS experiments using Ag/AgCl electrodes.

| subject | timepoint | N | impedance |  |  |  | offset |  |  |  |
| --- | --- | --- | --- | --- | --- | --- | --- | --- | --- | --- |
|  |  |  | mean | min | max | SD | mean | min | max | SD |
| human | pre Stim | 3 | 5.75 kΩ | 1.00 kΩ | 10.35 kΩ | 3.82 kΩ | 0.00 V | 0.00 V | 0.00 V | 0.00 V |
| human | post Stim | 3 | 5.42 kΩ | 9.55 kΩ | 9.55 kΩ | 3.50 kΩ | 0.00 V | 0.00 V | 0.10 V | 0.05 V |
| melon | pre Stim | 3 | 4.45 kΩ | 1.50 kΩ | 6.70 kΩ | 2.18 kΩ | 0.03 V | 0.00 V | 0.10 V | 0.05 V |
| melon | post Stim | 3 | 4.32 kΩ | 1.75 kΩ | 6.20 kΩ | 1.88 kΩ | 0.03 V | 0.00 V | 0.10 V | 0.05 V |

**Note:** The reported offset is rounded to one decimal place by the stimulation device. A reported offset of less than 0.1 V could therefore reflect a rounding error of a much smaller static offset (as the device is usually calibrated with an accepted offset in the range of millivolts) rather than an offset build up over time.

### Experimental setup

#### Human measurement, rubber electrodes

In both experiments, 10 minutes of EEG was recorded from 24 active sintered Ag-AgCl electrodes fitted in an elastic cap (EASYCAP GmbH, Herrsching, Germany) connected to an actiCHamp amplifier (Brain Products GmbH, Gilching, Germany) before and after 21 minutes of tACS at individual alpha frequency (2 mA peak-to-peak, 30 s fade-in and fade-out). Participants completed either vigilance or mental arithmetic tasks. EEG electrodes were placed according to the international 10-10 system, omitting the sites beneath the stimulation electrodes. tACS was delivered using carbonized rubber electrodes (7×5 cm over Cz and 5×5 cm over Oz), attached with a conductive paste (ten20, D.O. Weaver, Aurora, CO, United States). EEG electrodes were prepared with a conductive gel (SuperVisc, EASYCAP GmbH, Herrsching, Germany). The remote mode was started at the beginning of the pre-EEG block and switched off after the post-EEG block. Participants received either verum (all in experiment 1, 10 in experiment 2), or sham stimulation (10 in experiment 2, sham-protocol: 30 sec fade-in followed by 30 sec fade-out). Stimulation was applied at skin impedances of 10 kΩ or less.

### Human measurement, Ag/AgCl electrodes

For the human Ag/AgCl measurements, EEG was measured for 10 minutes before stimulation, 21 minutes during temporal interference stimulation and 10 minutes post stimulation from 28 active sintered Ag/AGCl-electrodes fitted in an elastic cap connected to an actiCHamp amplifier. EEG electrodes were placed according to the international 10-10 system, omitting the sites of the stimulation electrodes. During all blocks Participants completed a visual vigilance task. The stimulation was applied via two isolated neuroConn DC-STIMULATOR PLUS with extended frequency range (neuroConn GmbH, Germany) with one stimulator applying a sinusoidal current of 2 mA at 995 and the other stimulator applying a current of 2 mA at 1005 Hz. One stimulator was connected to two round sintered Ag/AgCl stimulation electrodes(Ø2 cm, neuroConn GmbH, Germany) situated at O1 and C3, the other stimulator was connected to electrodes at O2 and C4. Both pairs were fixed to the scalp using using Ten20-paste. All stimulation impedances were close to 10 kΩ or below.

### Melon measurement

A watermelon (*Citrullus lanatus*) of roughly Ø60-65 cm was used as a phantom head. Positions roughly corresponding to nosetip, FPz, Cz, Pz, P3, P4 and Oz were marked. EEG was recorded using an actiCHamp at 1000 Hz sampling rate without any pre-set filters with active sintered Ag-AgCl-electrodes. The reference electrode was placed on the nosetip, the ground at FPz and three recording electrodes on P3, P4 and Pz using adhesive rings and filled with Supervisc. The impedances were ensured to all be below 10 kΩ. For the carbonized rubber electrode test, a rectangular 7×5 cm rubber electrode was placed on the center of Cz and a second rectangular 5×5 cm electrode at the center of Oz and fixed to the melon using Ten20-paste. For the Ag/AgCl-test, two sintered Ag/AgCl round stimulation electrodes(Ø2 cm) were placed on the center of Cz and Oz and fixed using Ten20-paste. (The Ag/AgCl-recordings took place before the rubber-recordings to avoid any distortions to the surface area by Ten20-paste residue.) All stimulation impedances were below 10 kΩ.

The stimulation signal was generated in MATLAB as a perfect sine-wave of 10 Hz at 1000Hz sampling rate with zero-to-peak amplitude of 0.5 V and fed via a digital-to-analog converter (NI USB 6251, National Instruments, USA) to the remote port of a DC STIMULATOR PLUS, running in remote mode. A TTL-signal on the D/A-converter’s secondary analog output channel provided triggers for the EEG.

Each measurement started with a 10 minute EEG recording (stimulator active and in remote mode, D/A-converter set to 0 V), followed by 21 minutes of active tACS (with 30 seconds linear fade-in and fade-out), followed again by a 10 minute of EEG-recording (D/A-converter at 0 V).

For the recording of the square-wave responses, a square-wave at 10 Hz with a duty cycle of 20 % with an intercept of zero and an amplitude of 0.25 V and a duration of 1 minute was created in MATLAB and fed to the stimulator using the NI USB 6251. The second analog output channel of the D/A-conveter provided EEG-trigger information via TTL-pulses (same square-wave as stimulation but with an amplitude of 5V).

## References

Agrebi, F., Ghorbel, N., Ladhar, A., Bresson, S., & Kallel, A. (2017). Enhanced dielectric properties induced by loading cellulosic nanowhiskers in natural rubber: Modeling and analysis of electrode polarization. Materials Chemistry and Physics, 200, 155– 163.

Elyamany, O., Leicht, G., Herrmann, C. S., & Mulert, C. (2021). Transcranial alternating current stimulation (tACS): from basic mechanisms towards first applications in psychiatry. European Archives of Psychiatry and Clinical Neuroscience, 271 (1), 135–156.

Hampstead, B. M., Ehmann, M., & Rahman-Filipiak, A. (2020). Reliable use of silver chloride HD-tDCS electrodes. Brain Stimulation, 13 (4), 1005–1007.

Herrmann, C. S., Rach, S., Neuling, T., & Strüber, D. (2013). Transcranial alternating current stimulation: a review of the underlying mechanisms and modulation of cognitive processes. Frontiers in Human Neuroscience, 7, 279.

Langenbach, B. P., Savic, B., Baumgartner, T., & Knoch, D. (2020). Repeated anodal HDtDCS stimulation might render silver chloride electrodes unreliable. Brain Stimulation, 13 (3), 525–526.

von Conta, J., Kasten, F. H., Ćurčić-Blake, B., Schellhorn, K., & Herrmann, C. S. (2022). Characterizing low-frequency artifacts during transcranial temporal interference stimulation (ttis). Neuroimage: Reports, 2 (3), 100113.

Witkowski, M., Garcia-Cossio, E., Chander, B. S., Braun, C., Birbaumer, N., Robinson, S. E., & Soekadar, S. R. (2016). Mapping entrained brain oscillations during transcranial alternating current stimulation (tacs). Neuroimage, 140, 89–98.

Woods, A. J., Antal, A., Bikson, M., Boggio, P. S., Brunoni, A. R., Celnik, P., Cohen, L. G., Fregni, F., Herrmann, C. S., Kappenman, E. S., et al. (2016). A technical guide to tdcs, and related non-invasive brain stimulation tools. Clinical Neurophysiology, 127 (2), 1031–1048.

Zhu, X., & Deegan, T. (2024). Origin and mitigation of nonlinear artefacts in concurrent LFP and EEG recording [International meeting on temporal interference brain stimulation]. Retrieved June 23, 2026, from https://www.imperial.ac.uk/dementia-research-institute/events/international-meeting-on-temporal-interference-brain-stimulation/talk-recordings

